# Peripheral death by neglect and limited clonal deletion during physiologic B lymphocyte development

**DOI:** 10.1101/2023.05.30.542923

**Authors:** Mikala JoAnn Willett, Christopher McNees, Sukriti Sharma, Anna Minh Newen, Dylan Pfannenstiel, Thomas Moyer, David Stephany, Iyadh Douagi, Qiao Wang, Christian Thomas Mayer

## Abstract

Autoreactive B cells generated during B cell development are inactivated by clonal deletion, receptor editing or anergy. Up to 97% of immature B cells appear to die before completing maturation, but the anatomic sites and reasons underlying this massive cell loss are not fully understood. Here, we directly quantitated apoptosis and clonal deletion during physiologic B lymphocyte development using Rosa26^INDIA^ apoptosis indicator mice. Immature B cells displayed low levels of apoptosis in the bone marrow but started dying at high levels in the periphery upon release from bone marrow sinusoids into the blood circulation. Clonal deletion of self-reactive B cells was neither a major contributor to apoptosis in the bone marrow nor the periphery. Instead, most peripheral transitional 1 B cells did not encounter the signals required for positive selection into the mature B cell compartments. This study sheds new light on B cell development and suggests that receptor editing and/or anergy efficiently control most primary autoreactivity in mice.

**Summary:** The large amount of cell loss predicted during B lymphocyte development is unexplained by clonal deletion of self-reactive cells. Many transitional 1 B cells die in the periphery due to failed selection in the mature B cell compartments.

## Introduction

Antibody diversity is generated by V(D)J recombination during B cell development in the bone marrow and during somatic hypermutation in germinal centers. During B cell development, B cell-committed progenitors (pro-B cells; Fr. B/C) first recombine the immunoglobulin heavy chain (IgH) in the bone marrow parenchyma. Successful IgH rearrangement signals proliferation of large pre-B (Fr. CLJ) cells before resting small pre-B (Fr. D) cells begin VJ rearrangements at the immunoglobulin light chain (IgL) loci *Ig*κ and *Ig*λ (Hardy and Hayakawa, 2001). Immature (Fr. E) B cells expressing the fully rearranged surface IgM B cell receptor (BCR) leave the bone marrow via sinusoids and enter the blood circulation as transitional 1 (T1) B cells. In the spleen, signaling by the BCR and BAFF / BAFF-R are essential for B cell survival and differentiation into T2, mature follicular (FO) and marginal zone (MZ) B cells, a process also referred to as positive B cell selection (Loder et al., 1999; Sasaki et al., 2004; Schiemann et al., 2001; Turner et al., 1997).

Random BCR repertoire formation during B cell development inevitably creates the problem of self-reactivity. The clonal selection theory proposed that the period of antibody randomization is confined to a short time window during B cell development during which premature antigenic encounter results in “elimination” (clonal deletion) and self-tolerance instead of clonal expansion and antibody production (Burnet, 1959). Clonal deletion of immature B cells was first demonstrated in transgenic mice expressing a monoclonal self-reactive BCR (Nemazee and Burki, 1989). Alternative mechanisms were also discovered depending on the monoclonal BCR and the type / expression pattern of the self-antigen studied, which include silencing of the transgenic BCR by rearrangement of endogenous light chains (receptor editing) and anergy (Gay et al., 1993; Goodnow et al., 1988; Tiegs et al., 1993). Based on whether self-antigen is first encountered in the bone marrow or in the periphery, B cell tolerance mechanisms are categorized as central and peripheral, respectively. Subsequent work suggested that receptor editing rather than clonal deletion is the main mechanism of central B cell tolerance (Ait-Azzouzene et al., 2005; Halverson et al., 2004). To what extent clonal deletion contributes to peripheral tolerance is not entirely clear.

Up to 97% of newly developing immature B cells were estimated to die prior to becoming mature B cells (Allman et al., 1993; Rolink et al., 1998). If clonal deletion is rare during central tolerance, such a dramatic B cell loss could only be explained by either massive clonal deletion during peripheral tolerance or by high levels of cell death of immature B cells in the bone marrow and/or of peripheral transitional B cells by mechanisms other than clonal deletion of self-reactive B cells. The anatomic sites and mechanisms of cell loss during physiologic B cell development are therefore incompletely understood.

## Results

### Quantitation of apoptosis during physiologic B cell development

We analyzed Rosa26^INDIA^ mice by flow cytometry to quantitate apoptosis via FRET^+^ or FRET^neg^ fluorescent indicator states that depend on cleavage by active caspase-3 (aCasp3) and loss of FRET (Fig. 1A, fig. S1A, B)(Mayer et al., 2017). In the bone marrow, apoptosis was generally low in all B cell subsets examined, but significantly elevated in Fr. D pre-B cells (average 0.30% FRET^neg^, p<0.0001) and Fr. E immature B cells (average 0.14% FRET^neg^, p=0.016) compared with Fr. F mature recirculating B cells (average 0.09% FRET^neg^) (Fig. 1B, E). Apoptosis of earlier B cell precursors (Fr. B/C) was comparable to Fr. F. In contrast to the low levels of B cell apoptosis in the bone marrow, peripheral transitional 1 (T1) B cells underwent robust apoptosis in blood and spleen (average 1.5%-2% FRET^neg^; p<0.0001) compared with mature follicular (FO) B cells at these sites (average 0.16%-0.25% FRET^neg^; Fig. 1C-E). Slightly fewer splenic marginal zone (MZ) than FO B cells were FRET^neg^ (average 0.13%, p=0.016; Fig. 1D,E).

**Fig. 1.**
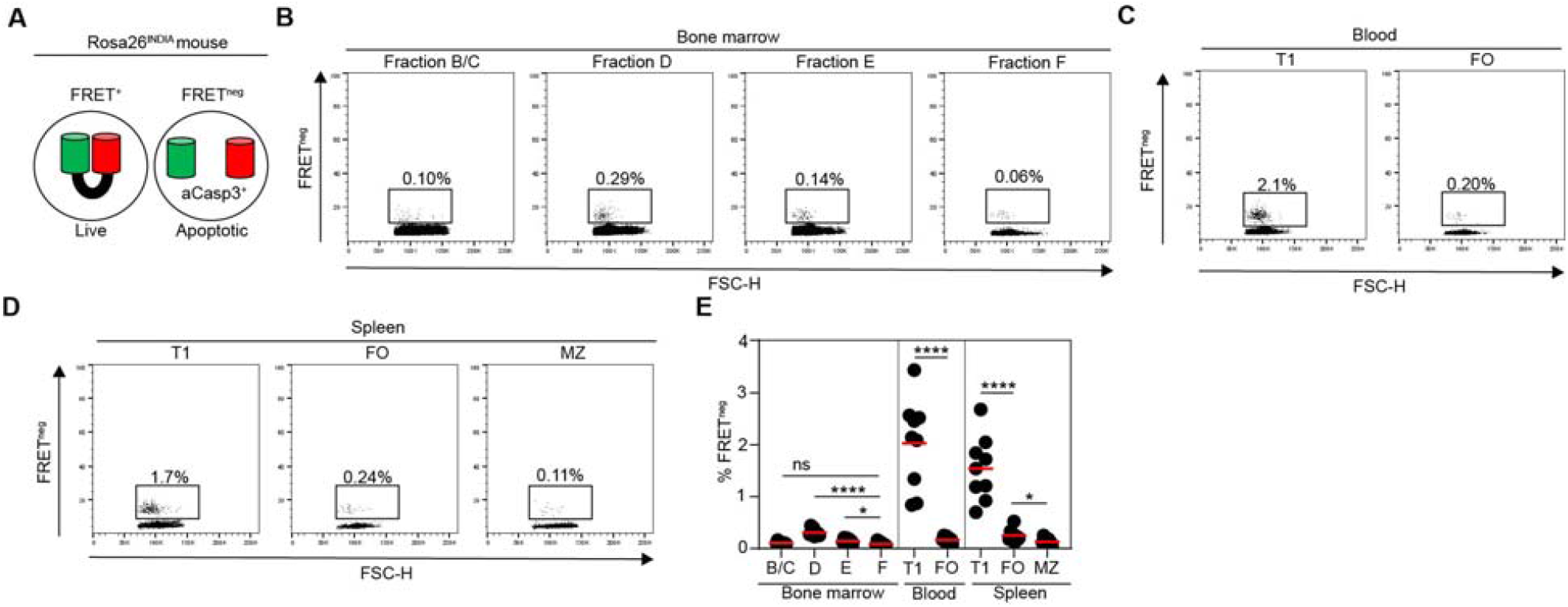
Quantitation of apoptosis during physiologic B cell development. (**A**) Schematics of Rosa26^INDIA^ apoptosis indicator mice. **(B-E)** Rosa26^INDIA^ mice were analyzed by flow cytometry (horizontal bars: mean values, B/C: pro/pre-B cells, D: small pre-B cells, E: immature B cells, F: mature recirculating B cells, T1: transitional 1 B cells, FO: mature follicular B cells, MZ: marginal zone B cells). **(B-D)** Representative dot plots depict FSC-H and FRET loss of indicated B cell developmental stages in (**B**) bone marrow, (**C**) blood and (**D**) spleen. (**E**) Quantitation of apoptotic FRET^neg^ cells. Data are combined from three independent experiments each involving three animals (**** p<0.0001, * p=0.016; unpaired student’s t-test).

We next used an independent staining panel to resolve T0, T1 and T2 transitional stages, as well as anergic T3 cells in blood and spleen (Henderson et al., 2010; Merrell et al., 2006). The least mature T0 cells died significantly more than T1 cells, and transitional B cells in the blood generally reached higher apoptosis levels than their counterparts in the spleen (fig.S1C). Splenic T2 cells returned to low levels of apoptosis (fig. S1C), suggesting that T2 differentiation and/or splenic localization are critical for B cell survival. Anergic T3 cells also exhibited low levels of apoptosis (fig. S1C).

We conclude that physiologic murine B cell development is associated with relatively low levels of apoptosis in the bone marrow and with high levels of apoptosis of T0/1 cells in the periphery.

### Immature B cell death upon release from bone marrow sinusoids

To distinguish immature B cells in the bone marrow parenchyma from those retained in bone marrow sinusoids prior to release into the blood circulation we performed intravascular labeling of Rosa26^INDIA^ mice (Pereira et al., 2009)(Fig.2A, B). FRET^neg^ cells were quantitated in bone marrow parenchyma, sinusoids, and the blood circulation. Immature B cells in bone marrow sinusoids and parenchyma displayed comparably low levels of apoptosis while blood T1 cells died at >10 times higher levels (Fig. 2C). Thus, release of immature B cells from bone marrow sinusoids into the blood circulation is associated with a dramatic increase in apoptosis.

**Fig. 2.**
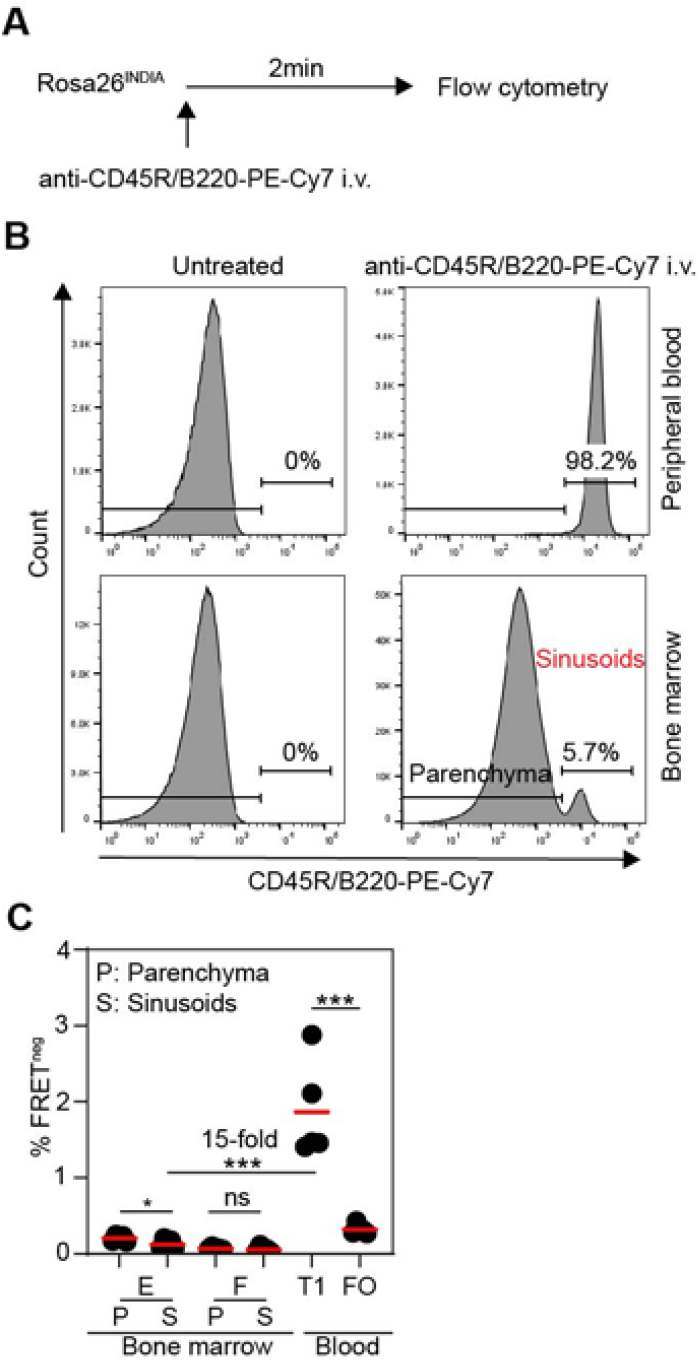
Immature B cell death upon release from bone marrow sinusoids into the blood circulation. **(A)** Schematics of the bone marrow sinusoid labeling experiment. **(B, C)** Rosa26^INDIA^ mice were injected intravenously with anti-CD45R/B220-PE-Cy7 two minutes prior to euthanasia. Untreated animals served as controls. **(B)** Representative histograms show PE-Cy7 labeling of CD19^+^Lineage(CD4.CD8α, NK1.1, F4/80, Ly-6G, Ter119)^neg^DAPI^neg^mRuby2^+^ B cells in either peripheral blood or bone marrow. All B cells in blood are labeled, as expected. Labeling in the bone marrow marks B cells in sinusoids. Non-injected mice show no labeling in either blood or bone marrow. (**C**) Quantitation of FRET^neg^ cells among Fr. E immature B cells and Fr. F mature recirculating B cells in the bone marrow parenchyma (P: non-labeled) or sinusoids (S: labeled). Labeled transitional 1 (T1) and mature follicular (FO) B cells in blood are shown as controls. Results are combined from three independent experiments each involving 1-2 animals (*** p<0.001, * p=0.042; unpaired student’s t-test).

### Limited clonal deletion of self-reactive B cells during physiologic B cell development

We next asked whether clonal deletion of self-reactive B cells contributes to cell loss during physiologic B cell development. We single cell-sorted FRET^+^ and FRET^neg^ Rosa26^INDIA^ B cell subsets, and then sequenced and cloned their BCR genes (Fig. 3A). Usage of downstream IgκJ4 and IgκJ5 elements is associated with receptor editing (Tiegs et al., 1993). IgκJ usage did not notably change in FRET^+^ B cells during development (fig. S2A). FRET^neg^ immature B cells used slightly less IgκJ5 (p=0.017) and more IgκJ1 than their FRET^+^ counterparts (fig. S2A), suggesting that some immature B cells die prior to excessive receptor editing. Autoreactive and polyreactive BCRs are enriched with positively charged amino acids in the IgH CDR3 region and with long IgH CDR3s (Mayer et al., 2020; Wardemann et al., 2003). The proportions of FRET^+^ B cells with ≥3 positive IgH CDR3 charges only modestly decreased during development (fig. S2B). FRET^neg^ early immature, immature and T1 B cell IgH CDR3 charges were not significantly different to FRET^+^ cells, yet more B cells with 4 positive charges were seen which were rare in any FRET^+^ B cell compartment (fig. S2B). The mean IgH CDR3 length did not notably change during B cell development, nor was there a significant difference in live and apoptotic B cells (fig. S2C).

**Fig. 3.**
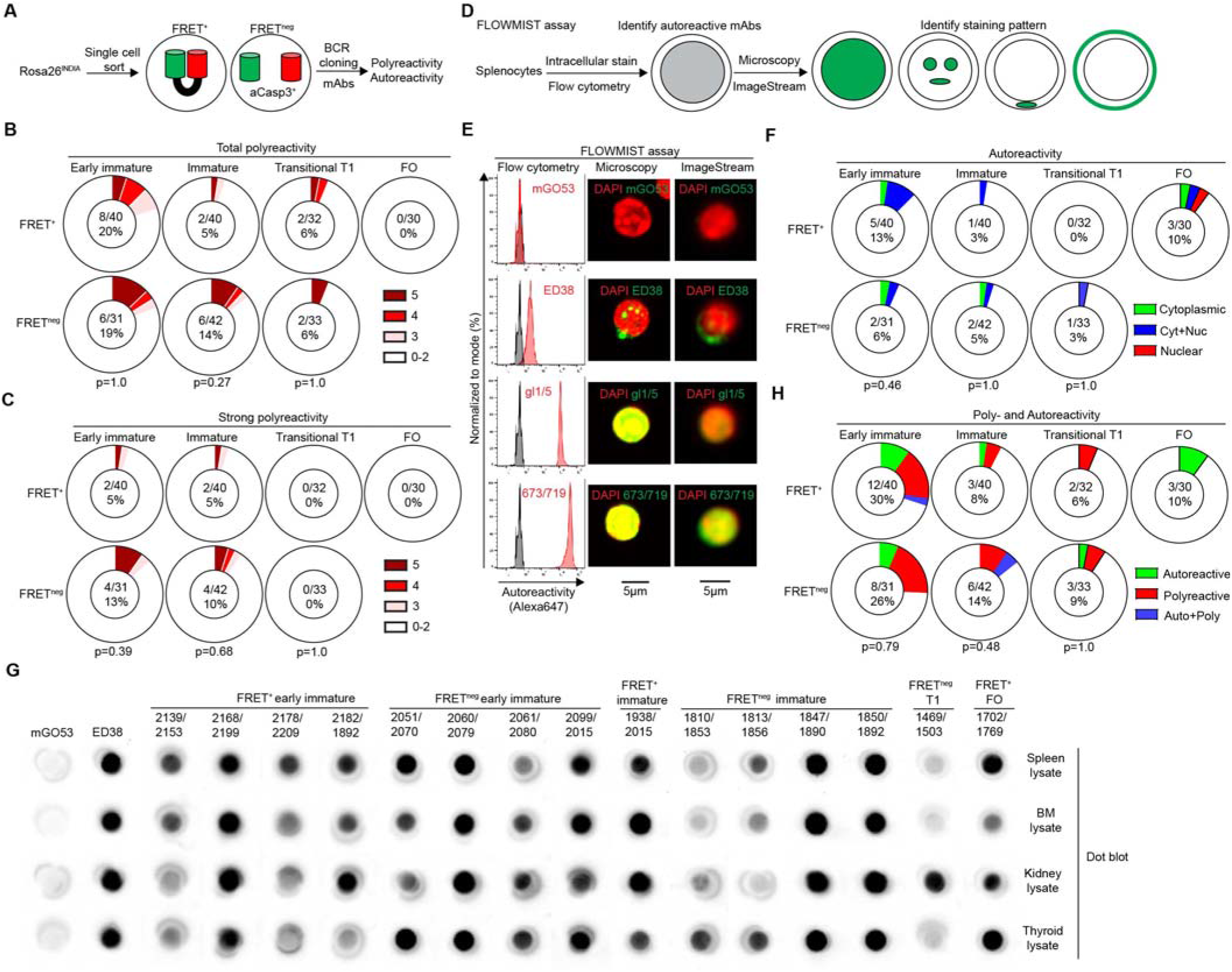
Limited clonal deletion of self-reactive B cells during physiologic B cell development. Live (FRET^+^) and apoptotic (FRET^neg^) Rosa26^INDIA^ B cell compartments were single-cell FACS-sorted. Ig genes were cloned and expressed as recombinant antibodies for testing. (**A**) Experimental outline. (**B, C**) Summary of polyreactivity determined by ELISA showing (**B**) total or (**C**) strong polyreactivity. Red intensity depicts polyreactivity to 3, 4 or 5 antigens, white represents reactivity to 2 or less antigens. (**D-F**) mAb autoreactivity testing by the FLOWMIST assay. (**D**) Experimental outline. (**E)** FLOWMIST analysis of control mAbs mGO53 (non-reactive), ED38 (cytoplasm/nucleus-reactive), 673/719 and its germline-revertant gl1/5 (both anti-nuclear). Left, flow cytometry histograms (unstained cells represented in grey). Right, fluorescent staining pattern (green) determined by confocal microscopy and ImageStream. DAPI is shown in red. Scale bars (5µm, bottom). (**F**) Summary of autoreactivity among B cells compartments. Green (anti-cytoplasmic), blue (anti-cytoplasmic and nuclear), red (anti-nuclear). (**G**) Dot blot analysis of mAb autoreactivity to C57Bl/6 spleen, bone marrow, kidney and thyroid lysates. Dot blots for mAbs reacting to at least one tissue are shown. mGO53 (non-reactive) and ED38 (highly polyreactive) antibodies were included as controls. Each reactive antibody was confirmed in two independent experiments. (**H**) Summary of autoreactivity (green) and polyreactivity (red) among indicated B cell compartments. Blue depicts antibodies that are autoreactive and polyreactive. (**A-H**) Pie chart centers show the number of reactive and total mAbs tested, and the percentage of reactive antibodies. The p value for comparing FRET^+^ and FRET^neg^ cells is shown below the pie charts (Fisher’s exact test). mAbs are derived from two to three independent single-cell sorts.

Since sequence features cannot accurately predict BCR binding properties, we next expressed 248 monoclonal antibodies (mAbs) corresponding to the BCRs of single FRET^+^ and FRET^neg^ B cells across development and directly tested polyreactivity and autoreactivity (Fig. 3A, Table S1). About 20% of FRET^+^ early immature bone marrow B cells were polyreactive, which dropped to 5-6% in FRET^+^ immature B cells and splenic T1 cells (Fig. 3B and fig. S2D). Polyreactivity further diminished in FRET^+^ FO B cells. Compared with FRET^+^ counterparts, FRET^neg^ B cells were not significantly enriched in polyreactivity in any compartment tested (Fig. 3B and fig. S2D, E). 5% of FRET^+^ early immature and immature B cells displayed strong polyreactivity, while only weakly polyreactive BCRs remained at the T1 stage (Fig. 3B, C and fig. S2D, E). Again, dying FRET^neg^ early immature and immature B cells did not contain significantly more polyreactive BCRs (Fig. 3B, C and fig. S2D, E).

We next developed a novel flow cytometry-based screening assay coupled with confocal microscopy or imaging flow cytometry (FLOWMIST) to identify anti-nuclear/subnuclear, cytoplasmic and/or membrane autoantibodies using primary mouse splenocytes as target cells (Fig. 3D). FLOWMIST sensitively and accurately detected autoreactive control antibodies previously tested by indirect immunofluorescence of HEp-2 cells (Fig. 3E) (Mayer et al., 2020; Meffre et al., 2004). After screening the 248 mAbs by FLOWMIST, we identified several weakly autoreactive antibodies recognizing cytoplasmic and/or nuclear antigens (fig. S2F, G). Some of these included highly polyreactive BCRs (see Table S1). Autoreactivity gradually declined from FRET^+^ early immature B cells to splenic T1 B cells but increased again in mature FO B cells (Fig. 3F, top), consistent with the finding that weak self-reactivity can promote positive selection into the mature FO B cell compartment (Noviski et al., 2019). Again, apoptosis had no detectable role at eliminating autoreactive B cells throughout B cell development (Fig. 3F, bottom; fig. S2F), similar to findings in germinal center B cells (Mayer et al., 2017; Mayer et al., 2020).

To exclude missed reactivity with fixation-sensitive epitopes and/or with tissue-specific self-antigens, we also performed immunoblot assays on all 248 antibodies using C57Bl/6 spleen, bone marrow, kidney, and thyroid lysates. All antibodies that consistently reacted to one or more tissue lysates had already been identified as either polyreactive and/or autoreactive (Fig. 3G, fig. S2H and Table S1).

To summarize, only small numbers of polyreactive and self-reactive B cells are eliminated by apoptosis throughout physiologic B cell development (Fig. 3H). At least 90% of the high level of cell death observed in peripheral T1 B cells cannot be attributed to self- or polyreactivity (Fig. 3H) and must therefore rely on other mechanisms.

### Death by neglect of T1 B cells

Interconnected signals transmitted through the BCR and BAFF-R are required for splenic T1 B cell development into T2 and mature B cells, a process also referred to as positive B cell selection (Rajewsky, 1996; Schiemann et al., 2001; Schweighoffer et al., 2013; Stadanlick et al., 2008; Turner et al., 1997). Considering that > 90% of the high level of T1 B cell apoptosis is not due to clonal deletion (Fig. 3), we hypothesized that most T1 B cells fail to receive positive selection signals and die by neglect. This would predict that T1 cells with lower IgM BCR expression are less competitive and die preferentially.

To this end, we analyzed B cell development in Nur77^GFP^ mice which report recent BCR signaling in splenic transitional and mature B cells by GFP expression (Zikherman et al., 2012). We found a small fraction of IgM^lo^GFP^neg^ cells among T1 B cells (average 3.5%), which gradually declined in T2/3 B cells (average 2%) and mature FO B cells (average 1.2%) (p<0.0001; Fig. 4A, upper panel; Fig. 4B, black symbols). Inhibition of B cell apoptosis in Nur77^GFP^Eµ-Bcl2^tg^ mice led to a marked accumulation of IgM^lo^GFP^neg^ T1 B cells (average 23%, p=0.0003, Fig. 4A, bottom panel; Fig. 4B, red symbols), indicating that T1 B cells with low BCR signaling normally die rapidly. Consistently, IgD^lo^IgM^lo^Fas^neg^ B cells in the blood and spleens of Rosa26^INDIA^ mice, which include IgM^lo^ T1 B cells, displayed markedly increased apoptosis compared with IgD^hi^IgM^+^ FO B cells (fig. S3). Despite inhibition of apoptosis, IgM^lo^GFP^neg^ cells were depleted from T2/3 B cells and mature FO B cells (average 2.9-3.4%), indicating that the BCR also controls T1 B cell differentiation independently from promoting survival (Fig. 4A, B)(Turner et al., 1997).

**Fig. 4.**
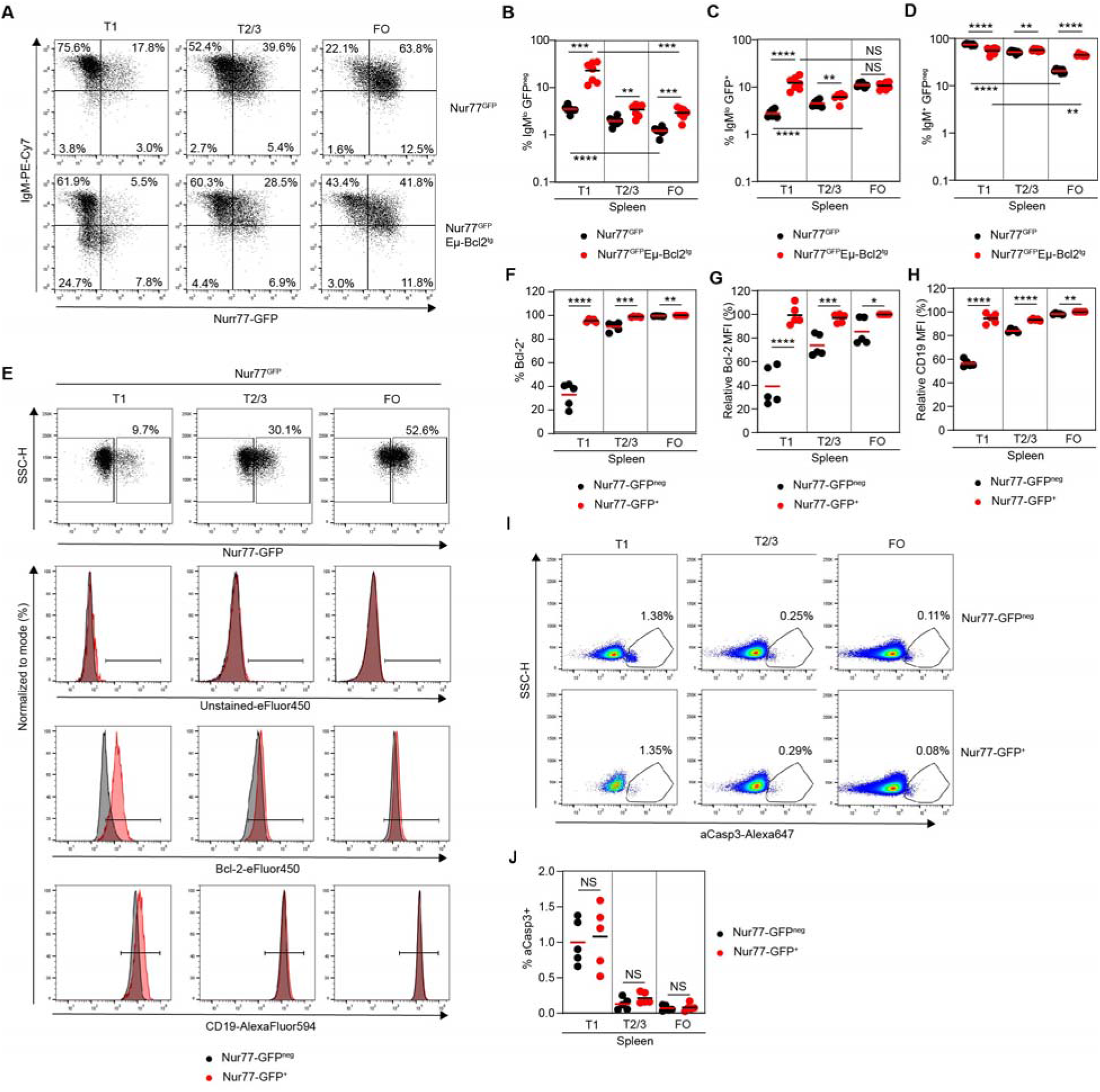
Peripheral death by neglect of T1 B cells. (**A-D**) Spleens of Nur77^GFP^ and Nur77^GFP^Eµ-Bcl2^tg^ mice were analyzed by flow cytometry. (**A**) Representative dot plots display GFP and IgM expression among indicated B cell subsets. (**B-D**) Quantitation of (**B**) IgM^lo^GFP^neg^, or (**C**) IgM^lo^GFP^+^, or (**D**) IgM^+^GFP^neg^ B cells (**** p<0.0001, *** p<0.0005, ** p<0.01; unpaired student’s t-test). Results are combined from three independent experiments each including 2-3 animals per genotype. (**E-J**) Spleen cells of Nur77^GFP^ mice were stained intracellularly for Bcl-2 and active caspase-3 (aCasp3) and analyzed by flow cytometry. GFP^+^ and GFP^neg^ subsets were compared. Results are from three independent experiments each including 1-2 animals. (**E**) Representative dot plots show GFP expression and SSC-H (top). Histograms (bottom) display intracellular staining for Bcl-2 and surface staining for CD19 (GFP^neg^: grey, GFP^+^: red). (**F-H**) Quantitation of (**F**) Bcl-2^+^ percentage or (**G**) Bcl-2 MFI or (**H**) CD19 MFI. MFIs are relative to GFP^+^ FO B cells. (**I**) Representative pseudocolor plots show aCasp3 vs SSC-H. (**J**) Quantitation of aCasp3^+^ cells among indicated subsets and genotypes (NS, not statistically significant; unpaired student’s t-test). (**A-J**) T1: transitional 1 B cells, T2/3: transitional 2/3 B cells, FO: mature follicular B cells, horizontal bars: mean values.

Nur77^GFP^EµBcl2^tg^ mice also accumulated IgM^lo^GFP^+^ T1 B cells compared with Nur77^GFP^ mice (p<0.0001; Fig. 4A and C) which could reflect autoreactive or polyreactive B cells that signaled and downregulated IgM (Zikherman et al., 2012). These differences were lost during subsequent B cell development (Fig. 4A, C). Additionally, IgM^+^GFP^neg^ B cells without recent BCR signaling gradually decreased during B cell development in Nur77^GFP^ mice but were forced to survive upon Bcl-2 expression and persisted significantly in the FO B cell compartment of Nur77^GFP^Eµ-Bcl2^tg^ mice (Fig. 4A and D).

To directly assess the relationship between BCR signaling and T1 B cell survival, we intracellularly stained Nur77^GFP^ B cells for Bcl-2 and active caspase-3 (aCasp3). While GFP^neg^ T1 B cells were Bcl-2^neg/lo^, almost all GFP^+^ T1 B cells expressed high levels of Bcl-2 comparable to mature B cells (Fig. 4E-G). In contrast, most GFP^neg^ T2/3 and FO B cells were Bcl-2^+^, and GFP^+^ counterparts only expressed slightly higher Bcl-2 levels (Fig. 4E-G). CD19, which is involved in BCR and BAFF-R signaling (Buhl et al., 1997; Jellusova et al., 2013), was also most differentially expressed at the T1 B cell stage as a function of GFP expression (Fig. 4E and H).

GFP^neg^ T1 B cells had a high level of aCasp3 staining (Fig. 4I,J), as expected (Fig. 1). Despite increased Bcl-2 and CD19 expression, GFP^+^ T1 B cells were not rescued from apoptosis (Fig. 4I, J). Thus, GFP^+^ T1 B cells in Nur77^GFP^ mice received recent survival signaling but lacked additional signals and/or T2 differentiation to complete positive selection. We conclude that T1 B cells compete for survival and differentiation in the periphery and that most of them die by neglect.

## Discussion

### Anatomic site of cell loss during B cell development

Calculations based on BrdU labeling experiments estimated that as little as 3% of immature B cells newly generated in the bone marrow enter the long-lived mature B cell pool in the spleen (Allman et al., 1993), predicting high levels of B cell loss by cell death. Additionally, most immature / transitional B cells were estimated to be lost in the bone marrow rather than the spleen (Allman et al., 1993; Rolink et al., 1998). Immature B cell loss in the bone marrow might be attributed to clonal deletion of self-reactive B cells, but several studies conflict with that hypothesis (Ait-Azzouzene et al., 2005; Halverson et al., 2004). It therefore remained incompletely understood at which anatomic sites and developmental stages B cells undergo apoptosis during physiologic B cell development, and why.

We here filled some of these gaps by directly measuring caspase-3-dependent apoptosis across B cell development using Rosa26^INDIA^ apoptosis indicator mice and by further discriminating T1 and T2 stages in the periphery. On average, only 0.14% of immature B cells in the bone marrow died, while this number increased to 2.0% of T1 cells in blood and to 1.5% of T1 cells in spleen. In these experiments, Rosa26^INDIA^ mice harbored on average 9.7 million total immature B cells in the bone marrow and 4.0 million total T1 cells in the spleen. We estimate the transitional B cell compartment in the blood at 0.20 million cells (2ml blood containing 5000 white blood cells per µl; 10% of white blood cells are B cells of which 20% are transitional cells). If 0.14% of immature B cells are lost every 20.6min (Mayer et al., 2017), assuming the speed of apoptosis is similar for all B cells, then approximately 0.95 million immature B cells would be lost per day in the bone marrow. Based on similar calculations and the observed fraction of dying cells, the peripheral T1 B cell compartments would be entirely eliminated within a day (0.28 Million T1 cells lost in the blood and 4.2 Million T1 cells in the spleen). While we cannot completely rule out that additional B cells are lost by a caspase-3-independent mechanism or that the measured fraction of apoptotic cells is underestimated for other reasons, our findings suggest that immature B cell loss by apoptosis in the bone marrow appears to be considerably lower than previously anticipated. One new angle provided in this study is that the peripheral blood circulation also needs to be considered as a potential site of cell loss which may in fact include the marginal zone sinuses and red pulp regions of the spleen from where circulating transitional B cells compete to enter the white pulp (Henderson et al., 2010). Indeed, we found that apoptosis returned to lower levels at the splenic T2 stage which coincides with the ability of T2 cells to enter B cell follicles (Loder et al., 1999). Our findings are consistent with a suppressive effect of the bone marrow microenvironment on immature B cell apoptosis (Sandel and Monroe, 1999) and with infrequent clonal deletion at this site (Ait-Azzouzene et al., 2005; Halverson et al., 2004). The factors negatively regulating B cell apoptosis in the bone marrow microenvironment are of interest for future investigation.

### Clonal deletion

One possibility for the massive B cell loss during B cell development is clonal deletion of autoreactive B cells as predicted by Burnet’s clonal selection theory (Burnet, 1957; Burnet, 1959). We directly quantitated clonal deletion across physiologic B cell development using Rosa26^INDIA^ apoptosis indicator mice, single-cell BCR cloning and extensive antibody profiling for self-reactivity and polyreactivity. Little clonal deletion was observed at the early immature and immature B cell stages in the bone marrow. This supports the view that other tolerance mechanisms such as receptor editing or anergy likely control most primary self-reactivity in C57Bl/6 mice (Ait-Azzouzene et al., 2005; Halverson et al., 2004; Zikherman et al., 2012). Importantly, we now also add evidence that peripheral T1 cells undergoing high levels of apoptosis were also not a major site of clonal deletion because most autoreactivity and polyreactivity had already been removed at this stage and >90% of the dying cells were not polyreactive or self-reactive.

While we cannot exclude that B cells reacting to self-antigens with such low affinity that they remain undetected in our assays are removed by apoptosis, this seems unlikely since apoptosis of developing B cells requires strong BCR stimulation by multivalent antigens (Hartley et al., 1991). Another possibility is that the cognate self-antigens were not present in our self-reactivity assays, which we minimized by comprehensively testing antibody binding to primary mouse cells, to various mouse tissue lysates and to defined self-antigens by using several independent methods rather than relying on the standard HEp-2 cell assay which is of human origin.

When comparing the amount of self-reactivity in FRET^+^ (live) B cell compartments in our study to other studies, one must consider that the mouse genetic background, environment, experimental models, methods, sensitivity and cut-off for detecting autoreactivity can differ among laboratories and the results may not be directly comparable. However, our finding that 8-30% of immature B cells in the bone marrow were autoreactive and/or polyreactive is consistent with the estimate that 25% of immature B cells undergo receptor editing, which is a known consequence of self-reactivity (Casellas et al., 2001). In Rosa26^INDIA^ mice, polyreactivity and autoreactivity decreased at the early immature to immature B cell stage, which is also seen in human B cell development (Wardemann et al., 2003). However, 30% self-/polyreactivity in mouse early immature B cells was considerably lower than the 55% to 75% observed in human early immature B cells (Wardemann et al., 2003). Many of the murine early immature B cells also showed relatively weak self-reactivity and polyreactivity. One possible explanation is that B cell development in C57Bl/6 mice generates less self-reactivity than B cell development in humans, possibly due to shorter average CDR3s in murine IgH chains (Shi et al., 2014), a feature known to associate with self- and polyreactivity (Mayer et al., 2020; Wardemann et al., 2003), or species differences in V(D)J segments. It is tempting to speculate that differences in early autoreactivity may also exist and influence autoimmune disease development in different mouse strains. For example, C57Bl/6 mice overexpressing Bcl-2 in B cells were found to be much less susceptible to autoimmune disease than the same mice on a mixed C57Bl/6 x SJL background (Strasser et al., 1992).

### Positive selection of T1 B cells

BCR signaling components (e.g. Syk, CD79a, CD45, CD19) and BAFF / BAFF-R control transitional B cell survival and differentiation into mature B cells, a process also referred to as positive selection that shapes the mature B cell repertoire (Cyster et al., 1996; Loder et al., 1999; Rickert et al., 1995; Sasaki et al., 2004; Schiemann et al., 2001; Turner et al., 1997). We found that during physiologic B cell development, only 10-20% of splenic T1 B cells exhibit recent BCR signaling. These cells expressed significantly higher levels of Bcl-2 and CD19 than their un-signaled counterparts, and resembled Bcl-2/CD19 expression of T2/3 cells and mature follicular B cells. Together with the significant drop in cell death from splenic T1 to T2 cells and the scarcity of clonal deletion, we hypothesize that the majority of T1 cells fail positive selection and undergo death by neglect. A recent study by Hobeika and colleagues supports this view because *in vivo* stimulation of T1 B cells by anti-IgM antibodies did not induce apoptosis as seen *in vitro*, but instead caused Bcl-2 expression, survival and developmental progression into T2 cells (Hobeika et al., 2018). T1 cell death by lack of survival signaling and positive selection resembles the mechanism of cell death in the light zones of germinal centers (Chen et al., 2023; Mayer et al., 2017).

The process of T1 B cell positive selection warrants further investigation. It seems unlikely that positive selection solely reflects self-antigen-triggered BCR signaling since 80% of mature follicular B cells experience ongoing signaling as indicated by the Nur77^GFP^ reporter, whereas only up to 10% are detectably autoreactive and/or polyreactive. One possibility is that the signaling threshold to induce Nur77^GFP^ is much lower than the detection threshold of our self-reactivity assays. Another point to consider is that the BCR and BAFF-R receptor systems were shown to interact and even share signaling components (Jellusova et al., 2013; Schweighoffer et al., 2013; Stadanlick et al., 2008). It is therefore possible that the Nur77^GFP^ reporter indicates processes in addition to antigen triggered BCR signaling *in vivo*. Nevertheless, BCR-dependent positive selection predicts that weakly autoreactive T1 B cells have an advantage in positive selection, and this is what we and others observed (Noviski et al., 2019).

Open questions also remain about the purpose of positive selection during peripheral B cell development at the expense of substantial cell loss. It is tempting to speculate that advantageous features are being selected and/or that detrimental properties other than autoreactivity/polyreactivity are being avoided in the repertoire.

## Materials and Methods

### Mice and treatments

Animals were maintained under specific pathogen-free conditions at NCI (Frederick and Bethesda). For bone marrow sinusoid B cell labeling, mice were injected intravenously with 2µg PE-Cy7-conjugated anti-CD45R/B220 antibody (Thermo Fisher Scientific, 25-0452-82) 2 min prior to euthanasia (Pereira et al., 2009). All animal procedures reported in this study that were performed by NCI-CCR affiliated staff were approved by the NCI Animal Care and Use Committee (ACUC) and in accordance with federal regulatory requirements and standards. All components of the intramural NIH ACU program are accredited by AAALAC International.

### Enzyme-linked immunosorbent assay (ELISA)

Polyreactivity was assessed by measuring antibody binding to LPS (Sigma, L2637), Keyhole Limpet Hemocyanin (KLH) (Sigma, H8283-100 MG), double stranded DNA (dsDNA) (Sigma, D4522), single stranded DNA (ssDNA, prepared from dsDNA) and human insulin (Sigma, I9278) by ELISA as described (Gitlin et al., 2016). Monoclonal antibodies and control antibodies (mGO53, non-reactive; ED38, highly polyreactive) containing human IgG1 constant regions were tested at 4µg/ml and at three four-fold dilutions and were detected with peroxidase-conjugated goat anti-human IgG Fc (Jackson ImmunoResearch, 109-035-098). Peroxidase was revealed with 1-Step ABTS substrate solution (Thermo Fisher, 37615) for 30 min and absorbance at 405nm was measured with a SpectraMax iD3 Multi-Mode Microplate Reader (Molecular Devices). A_405_ values were corrected for PBS-only signal for each plate. Monoclonal antibodies binding to ≥3 different antigens at 4µg/ml above a defined threshold were considered polyreactive. Those antibodies still binding to ≥3 different antigens at 1µg/ml were considered strongly polyreactive.

### Flow cytometry and cell sorting

FACS buffer (PBS containing 0.1% w/v BSA and 2mM EDTA) was used for cell isolation, incubation and acquisition if not stated otherwise. For Rosa26^INDIA^ experiments, euthanized mice, buffers, and consumables contacting cells were pre-chilled and maintained on ice to minimize *de novo* apoptosis during cell isolation, staining and acquisition. All centrifugations and cell sorting were done at 4°C.

Immediately after euthanasia peripheral blood was drawn from the inferior vena cava, mixed with 10mM EDTA to prevent coagulation and added to 10ml ACK lysis buffer (Quality Biological, 118-156-101). After incubation for 5 min on ice, cells were centrifuged for 7 min at 1300 RPM and washed with 10ml of FACS buffer. Bone marrow was flushed out from femurs and tibiae. Single cell suspensions were created from bone marrow and spleens by forcing the tissue through 70 µm cell strainers (Corning, 352350). For spleen, erythrocytes were lysed by resuspending pellets in 1ml ACK lysis buffer (Quality Biological, 118-156-101) and incubating for 1 min on ice, followed by washing with FACS buffer. Blood leukocytes, bone marrow and spleen cell suspensions were then transferred to 96-well round bottom plates. All centrifugations were performed for 3 min at 1300 RPM.

For live cell staining, Fc receptors were first blocked for 15 min on ice with rat anti-mouse CD16/CD32 antibody (2.4G2, produced and purified by the EIB Flow Cytometry Core). After centrifugation, cells were stained with antibodies to surface antigens for 45 min at 4°C and washed three times with FACS buffer. If applicable, cells were then stained with fluorescently conjugated streptavidin for 10 min at 4°C and washed three times. Cells were resuspended in FACS buffer containing 0.2µg/ml propidium iodide (PI) (Sigma-Aldrich, P4170) or 0.1µg/ml DAPI (Sigma-Aldrich, D9542) prior to acquisition to exclude dead/necrotic cells.

For intracellular staining and for some live cell experiments where PI and DAPI were not compatible with the staining panel, cells were washed twice with PBS and stained using the Zombie NIR Fixable Viability Kit (Biolegend, 423106, diluted 1:500 in PBS) for 30 min at 4°C prior to Fc blocking and staining of surface antigens. For intracellular staining, cells were then washed with PBS and incubated with Fixation/Permeabilization solution (BD Biosciences, 51-2090KZ) for 30 min at 4°C. Cells were washed two times with Perm/Wash buffer (BD Biosciences, 51-2091KZ) and then incubated with antibodies against intracellular antigens diluted in Perm/Wash buffer for 45 min at 4°C. After three washes with Perm/Wash buffer, cells were resuspended in FACS buffer.

Cells were acquired on LSRII and LSRFortessa flow cytometers (BD Biosciences). Rosa26^INDIA^ cells were acquired on LSRFortessa and FACSymphony A5 flow cytometers (BD Biosciences) with the following specifications: mNeonGreen (488nm excitation, 530/30 BP, 505 LP), FRET (488nm excitation, 610/20 BP, 600 LP), mRuby2 (561nm excitation, LSRFortessa: 582/15 BP; FACSymphony: 585/15 BP, 570 LP). Rosa26^INDIA^ bulk cell sorting was performed on FACSAria and Fusion instruments (BD Biosciences) with the same specifications indicated above for LSRFortessa. Flow cytometry data were analyzed using FlowJo software versions 10.7 or 10.8 (BD Biosciences). Doublets were excluded using FSC-W. For Rosa26^INDIA^ analysis and cell sorting, FRET^loss^ was derived as new parameter by dividing mNeonGreen and FRET signals, as done previously (Mayer et al., 2017).

Gating for Rosa26^INDIA^ experiments was performed as follows:

In Figure 1, bone marrow B cells were gated CD19^+^mRuby2^+^DAPI^neg^Lineage(CD4, CD8α, F4/80, NK1.1)^neg^ and CD43^+^IgD^neg^IgM^neg^ (Fr. B/C), CD43^neg^IgD^neg^IgM^neg^ (Fr. D), CD43^neg^IgD^neg/lo^IgM^+^ (Fr. E) or CD43^neg^IgD^hi^IgM^+^ (Fr. F). Blood and spleen B cells were gated CD19^+^mRuby2^+^DAPI^neg^Lineage(CD4, CD8α, F4/80, NK1.1, CD95)^neg^ and IgD^lo^IgM^hi^CD21^neg^ (transitional 1, T1), IgD^lo^IgM^hi^CD21^hi^ (marginal zone, MZ) or IgD^hi^IgM^+^ (follicular, FO).

Gating in Figure 2 was performed as in Figure 1 except intravascular anti-CD45R/B220-PE-Cy7 was used to label bone marrow sinusoids, Lineage additionally included Ter-119 and Ly-6G markers and dead cells were excluded using Zombie-NIR.

Blood and spleen B cells in Figure S1C were gated CD19^+^mRuby2^+^DAPI^neg^Lineage(CD4, CD8α, F4/80, NK1.1, Ly-6G, Ter-119, CD95)^neg^ and AA4.1^+^IgD^neg^CD23^neg^IgM^+^ (transitional 0 B cells, T0), AA4.1^+^IgD^+^CD23^neg^IgM^+^ (transitional 1 B cells, T1), AA4.1^+^IgD^+^CD23^+^IgM^hi^ (transitional 2 B cells, T2), AA4.1^+^IgD^hi^CD23^+^IgM^lo^ (anergic B cells, T3), AA4.1^neg^IgD^+^CD23^+^IgM^+^CD21^+^ (mature follicular B cells, FO) and AA4.1^neg^IgD^low^CD23^low^IgM^hi^CD21^hi^ (marginal zone B cells, MZ).

Gating in Figure S3 was CD19^+^mRuby2^+^DAPI^neg^Lineage(CD4, CD8α, F4/80, NK1.1, CD95)^neg^ and IgD^lo^IgM^lo^Igλ^neg^ (BCR^lo^) or IgD^hi^IgM^+^ (follicular, FO).

For Rosa26^INDIA^ bulk sorting in Fig.S1B, the following gating was done on bone marrow and spleen cells: B220^+^Lineage(CD4, CD8α, NK1.1, F4/80, Ly-6G, Ter-119)^neg^DAPI^neg^mRuby2^+^ and FRET^+^ or FRET^neg^. In case of splenic B cells, GL7^hi^ germinal center (GC) B cells were additionally excluded.

Gating for other experiments was performed as follows:

In Figure 4, spleen B cells were gated PI^neg^CD138^lo^TACI^lo^CD19^+^Fas^neg^ (Figure 4A-D) or Zombie-NIR^neg^Lin(CD4,CD8α,NK1.1,Ly-6G,Ter-119,F4/80)^neg^CD19^+^Fas^neg^ (Figure 4E-J) and AA4.1^+^CD21^neg^CD23^neg^ (transitional 1 B cells, T1), AA4.1^+^CD21^lo^CD23^+^ (transitional 2/3 B cells, T2/3) or AA4.1^neg^GL7^neg^CD21^+^CD23^+^ (mature follicular B cells, FO).

### Single B cell sorting, antibody cloning and recombinant expression

Euthanized Rosa26^INDIA^ mice, buffers, and consumables contacting cells were pre-chilled and maintained on ice to minimize *de novo* apoptosis during cell isolation, staining and cell sorting. Centrifugation and cell sorting were done at 4°C. Typically, two mice were pooled per sort. Staining was performed as described in the “Flow cytometry” section. Bone marrow cells were gated B220^+^Lineage(CD4, CD8α, NK1.1, F4/80)^neg^Zombie-NIR^neg^mRuby2^+^CD43^neg^ and IgM^neg^IgD^neg^ (Fr. D) or IgM^+^IgD^neg/lo^ (Fr. E). For one Fr. D sort, bone marrow cells were sorted separately from two individual mice, dead cells were excluded with DAPI and Lineage additionally included Ly-6G/Ly-6C and Ter-119. B lymphocytes were enriched from erythrocyte-lysed spleen cells using CD43 MicroBeads (Miltenyi Biotec,130-049-801) and LS columns (Miltenyi Biotec, 130-042-401) according to the manufacturer’s instructions prior to staining. Spleen cells were gated CD19^+^Lineage(CD4, CD8α, NK1.1, F4/80)^neg^Zombie-NIR^neg^mRuby2^+^Fas^neg^GL7^lo^ and IgD^lo^IgM^hi^CD21^lo^ (transitional 1, T1) or IgD^hi^IgM^+^ (mature follicular B cells, FO). For each B cell population, FRET^+^ and FRET^neg^ single B cells were sorted into 96-well PCR plates containing 4µl lysis buffer (for composition see (von Boehmer et al., 2016)) and were immediately frozen on dry ice. Plates were directly processed or stored at - 80°C. cDNA preparation, Ig gene amplification, sequencing and cloning into human expression vectors, recombinant expression and antibody purification were essentially done as described (Mayer et al., 2017; Mayer et al., 2020; von Boehmer et al., 2016) except that only IgM-specific reverse primers were used for IgH amplification, and optimized primers were used for Igκ and Igλ sequencing after the 2^nd^ PCR (for a list of all primers used see Table S2). Ig sequences were analyzed with IMGT/V-QUEST and IgBlast and were directly synthesized for cloning (IDT). Fr. D cells carrying a functional BCR were considered early immature B cells (Wardemann et al., 2003). 293-F cells (Thermo Fisher, R79007) were maintained in 293 Freestyle medium (Thermo Fisher, 12338026) and were transiently co-transfected with IgH and IgL expression vectors for seven days prior to monoclonal antibody purification from the culture supernatant using Protein G Sepharose 4 Fast Flow beads (Cytiva,17-0618-05) and IgG elution buffer (Thermo Fisher, 21009). After neutralization and buffer exchange to PBS, the IgG concentration was determined using NanoDrop One (Thermo Fisher, ND-ONE-W). All monoclonal antibodies were assessed for integrity by SDS-PAGE analysis.

### FLOWMIST assay for autoreactivity

Spleen cells were prepared for intracellular staining as described under “Flow cytometry and cell sorting” with the following modifications: cells were stained with the Live/Dead Fixable Aqua Dead Cell Stain Kit (Invitrogen, L34966) for 30 min at 4°C (1:500 dilution in PBS) prior to Fc blocking. Surface staining with anti-CD3ε-PE (hamster IgG, not recognized by the secondary antibody used in later steps) was additionally performed. Cells were fixed using the Foxp3/Transcription Factor Staining Buffer concentrate and diluent (Thermo Fisher Scientific, 00-5523-00) for 30 min at 4°C. Fixed cells were washed two times with 1x perm/wash buffer (BD, 554723) or with 1x Permeabilization Buffer (Thermo Fisher Scientific, 00-8333-56). Monoclonal antibodies were diluted to 10µg/ml in 1x perm/wash buffer or 1x Permeabilization Buffer and cells were incubated for 1 hour at 4°C followed by three wash steps. Bound autoreactive antibodies were detected with 1µg/ml AlexaFluor647-conjugated F(ab’)2 goat anti-human IgG-Fc (Jackson ImmunoResearch, 109-606-098). Secondary staining was performed in same manner as primary staining. After another three washes, cells were resuspended in FACS buffer and acquired on a BD LSRFortessa X-20 Cell Analyzer (BD Biosciences). Mean AlexaFluor647 fluorescence intensity was calculated for Aqua^neg^CD3ε ^+^ gated T cells. The MFI ratio was calculated relative to mGO53 negative control monoclonal antibody. Monoclonal antibodies were deemed autoreactive if the MFI ratio was greater than 3.

Autoreactive monoclonal antibodies were further analyzed for the fluorescence pattern as follows. Intracellular staining was performed as described above but without Aqua staining. After secondary antibody incubation, cells were stained with 5µg/ml DAPI for 5 min at room temperature prior to washing three times. For imaging flow cytometry analysis, cells were resuspended in FACS buffer and acquired on Amnis ImageStream^x^ MKII using INSPIRE (v200.1.681.0) software at 60x magnification. Images were analyzed using IDEAS v 6.3 software and standard ImageStream analysis wizards and tools/algorithms. For confocal microscopy, cells were resuspended in PBS and transferred to a 35 mm dish with glass coverslip (Ibidi, 81156). Cells were incubated at room temperature for 10 min, then PBS was aspirated and 400µl fresh PBS were added to cover cells. Cells were imaged with 60x magnification on a Nikon Ti2 / Yokogawa CSU-W1 spinning disk confocal microscope with a Hamamatsu Orca Flash 4.0 camera. Confocal images were analyzed using ImageJ.

### Dot blot assay for autoreactivity

The dot blot assay was adapted from (Ahrens et al., 2012). The following lysis buffer was prepared freshly and kept on ice: 1xTBS (Thermo Fisher, 28358) supplemented with 1% NP-40 (Sigma, I8896), Complete Protease Inhibitor Cocktail (Roche, 11873580001) and 1x Halt Phosphatase Inhibitor Cocktail (Thermo Fisher, 78428). Spleen, kidney and thyroid of C57Bl/6 mice were cut into small pieces. About 100mg tissue was frozen in liquid nitrogen, then placed on ice and submerged in 4ml lysis buffer. Bone marrow was directly flushed out from 2 femurs and 2 tibiae with 4ml lysis buffer. Tissue was homogenized on ice for 1 min on medium setting using a Fisherbrand 150 Homogenizer (Fisher Scientific, 15-340-167) and Plastic disposable generator probes (Fisher Scientific, 15-340-176). Lysates were agitated for 2 hours on ice, centrifuged for 20 min at 14,000 RPM and the supernatant was harvested and stored at -80°C. Protein concentrations were determined with the Pierce BCA Protein Assay Kit (Thermo Fisher, 23225) according to the manufacturer’s instructions. Dot blots were performed using Nitrocellulose membranes (Thermo Fisher, 88018) and the Bio-Dot Microfiltration Apparatus (Bio Rad, 1706545) according to the manufacturer’s instructions. In brief, 1µg lysate was loaded per well in 1xTBS, and wells were then blocked for 2 hours at room temperature using 2% BSA in 1xTBS. After washing with 1xTBS 0.05% Tween20, monoclonal antibodies were incubated at 10µg/ml for 2h at room temperature. After washing, secondary peroxidase-conjugated goat anti-human IgG Fc (Jackson ImmunoResearch, 109-035-098) was incubated at room temperature for 1h at 1:5000 dilution in blocking buffer containing 0.05% Tween20. After washing, the membrane was removed from the apparatus, washed in 1xTBS and then incubated for 5 min with SuperSignal West Pico PLUS Chemiluminescent Substrate (Thermo Fisher, 34577). The chemiluminescent signal was scanned with UVP ChemStudio (Analytik Jena). Images were analyzed with ImageJ.

### Statistical analysis

Statistical significance was determined using GraphPad Prism software. Data were first evaluated for normal distribution by Anderson-Darling test, D’Agostino & Pearson test Shapiro-Wilk test and Kolmogorov-Smirnov test. If any test reported N is too small for evaluation or if data were normally distributed, then unpaired student’s t-test was used. If data did not pass normality tests, Mann-Whitney test was used. Pie charts were compared using Fisher’s exact test. Test results are indicated in the Figures and Figure legends.

## Supporting information

Auxiliary Table S1

Auxiliary Table S2

## Acknowledgments

We thank all members of the Experimental Immunology Branch, particularly Alfred Singer, Hyun Park, Richard Hodes, Vanja Lazarevic, Paul Roche, Susan Sharrow for critical discussions and advice; Jeffrey Chiang for technical support; Assiatu Crossman, Larry Granger, Tony Adams, William Hajjar, Tengfei Zhang for flow cytometry support; Jan Wisniewski for microscopy support. We thank Jagan Muppidi for discussions and for kindly providing mice, and all staff at the NCI Frederick and NCI Bethesda animal facilities for their critical help, particularly Jennifer Wise. This work was supported by the Intramural Research Program of the National Cancer Institute, Center for Cancer Research, National Institutes of Health. This research was supported in part by the Intramural Research Program of the NIH, National Institute of Allergy and Infectious Diseases. C.T.M. is a Stadtman Investigator. M.J.W., S.S., A.M.N. and D.P. are CRTA fellows. The authors declare no conflicts of interests.

## Supplementary Materials

Figs. S1 to S3

Auxiliary table legends for tables S1 to S2

Auxiliary Tables S1 to S2

