## Supplementary Figures for "Peripheral death by neglect and limited clonal deletion during physiologic B lymphocyte development"

Mikala JoAnn Willett, Christopher McNees, Sukriti Sharma, Anna Minh Newen, Dylan Pfannenstiel, Thomas Moyer, David Stephany, Iyadh Douagi, Qiao Wang

and Christian Thomas Mayer

**This PDF file includes:**

Figs. S1 to S3

Legends for auxiliary tables S1 to S2

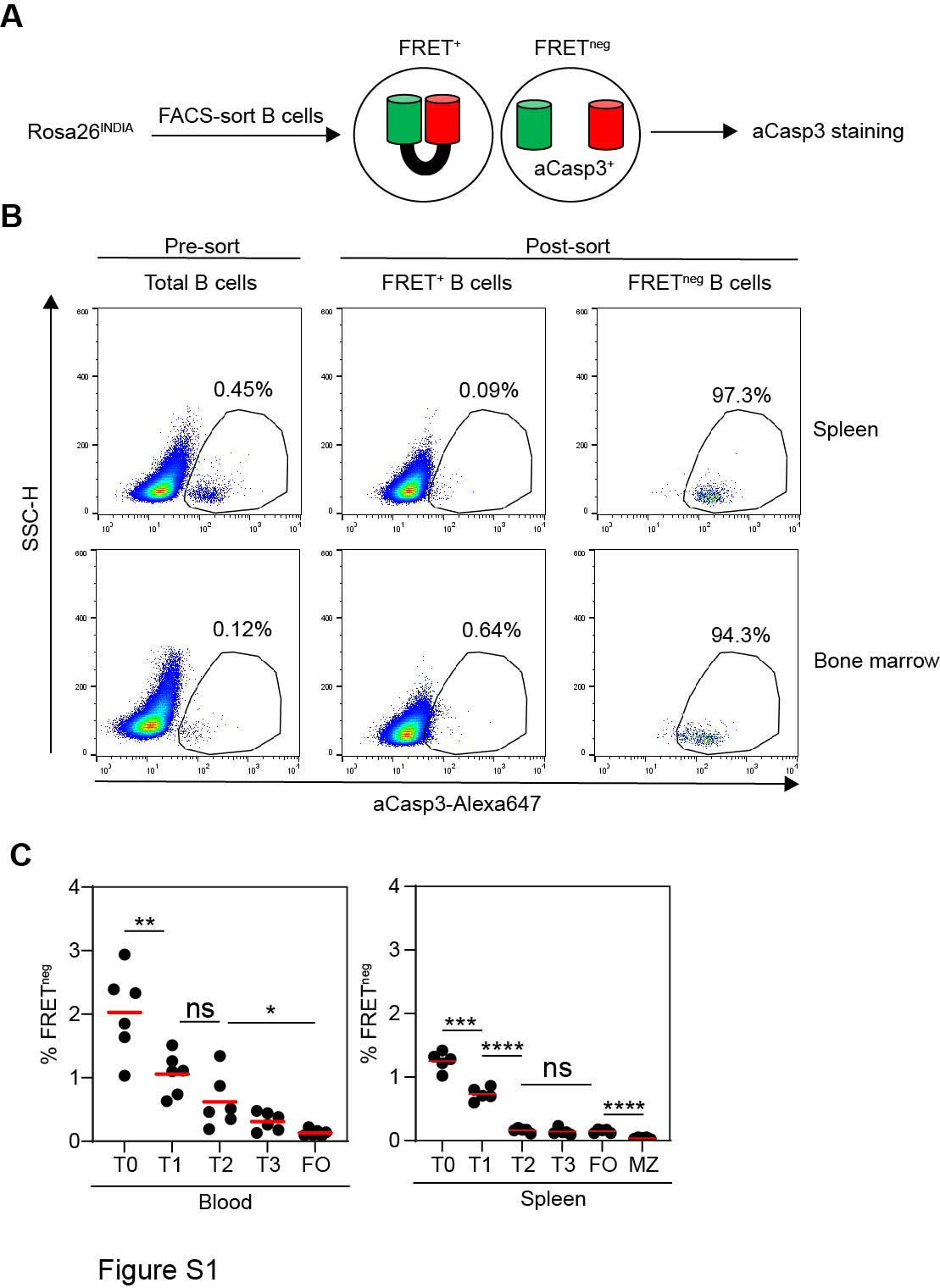

**Fig. S1. Quantitation of apoptosis during physiologic B cell development.**

**(A)** Schematics of the FACS sorting experiment. **(B)** FRET^+^ and FRET^neg^ B cells were FACS-sorted from bone marrow and spleen of Rosa26^INDIA^ mice. Immediately after FACS-sorting, dead/necrotic cells were stained with Zombie-NIR, followed by intracellular staining of early apoptotic cells with aCasp3. Pre-sort B cells served as control. aCasp3 staining and SSC-H is shown for indicated Zombie-NIR^neg^ B cell populations before and after sorting. Percentages of aCasp3^+^ cells are shown. One of two independent experiments is shown. **(C)** Blood and spleen B cell subsets of Rosa26^INDIA^ mice were analyzed by flow cytometry. FRET loss is quantitated in the indicated B cell subsets (T0, T1, T2: transitional B cells, T3: anergic B cells, FO: mature follicular B cells, MZ: marginal zone B cells). Horizontal bars: mean values. Results are combined from three independent experiments each including 1-2 animals (**** p<0.0001, *** p=0.0002, ** p=0.0095, * p=0.0179; unpaired student’s t-test).

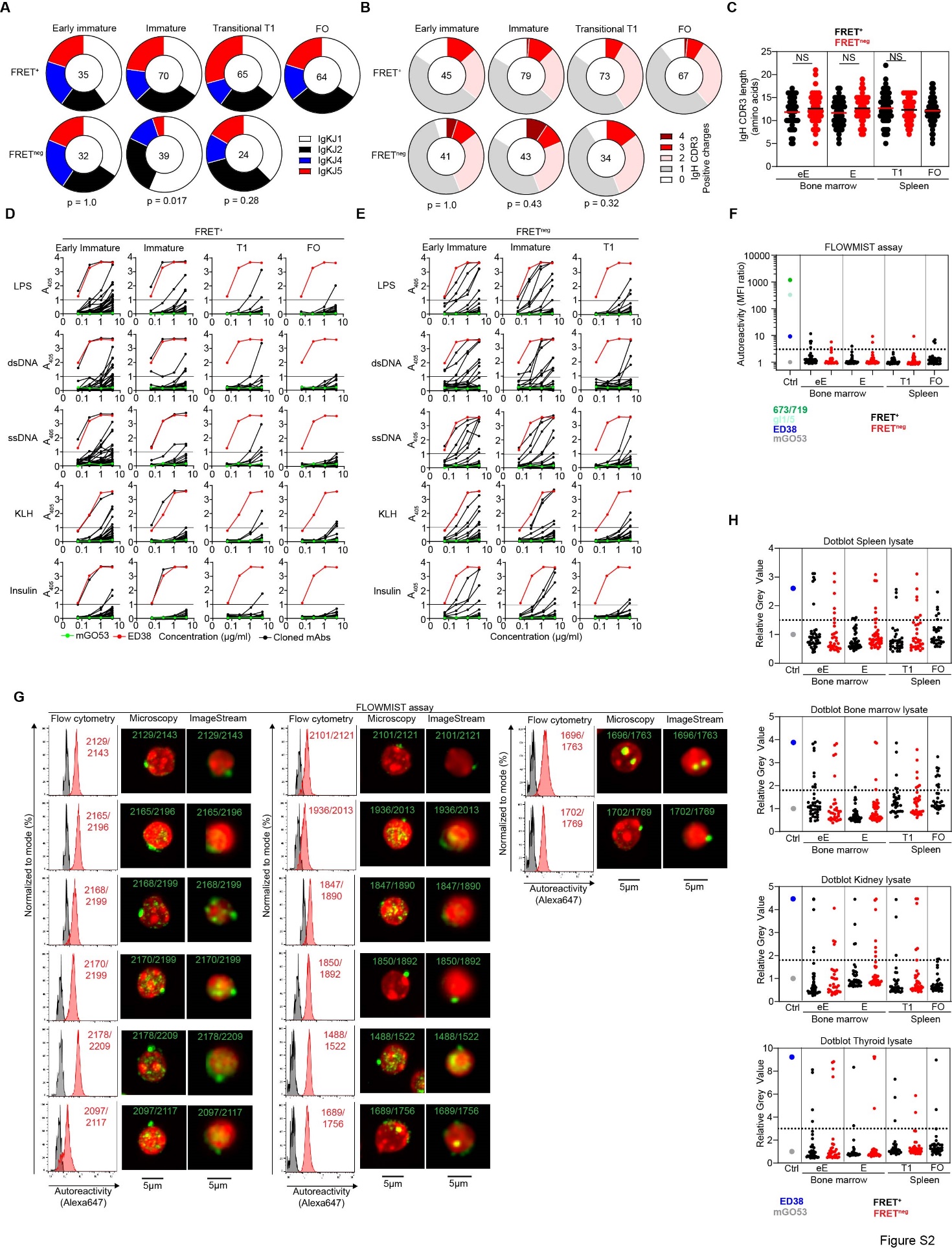

**Fig. S2. Limited clonal deletion of autoreactive and polyreactive B cells during physiologic B cell development.**

Live (FRET^+^) and apoptotic (FRET^neg^) B cells of various developmental stages were single-cell FACS-sorted from the bone marrow (eE: early immature, E: immature) and spleen (T1: transitional 1, FO: mature follicular) of Rosa26^INDIA^ mice. Ig genes were PCR amplified, sequenced, cloned into expression vectors and the corresponding recombinant antibodies were expressed, purified, and tested. (**A-C**) Ig sequence analysis of indicated B cell compartments. Only cells with paired functional heavy- and light chains were considered for analysis. **(A)** IgκJ usage. **(B)** Number of positively charged amino acids in the IgH CDR3 region. (**A, B**) Numbers in the center of each pie indicate the numbers of BCRs assessed. The p value for comparing IgκJ5 and IgH CDR3s with at least three positive charges among FRET^+^ and FRET^neg^ cells are shown below the pie charts (Fisher’s exact test). **(C)** IgH CDR3 lengths. Each dot represents one cloned BCR. Horizontal bars: mean values (NS, not statistically significant; Mann-Whitney test. **(D, E)** ELISA measurements show binding of monoclonal antibodies cloned from the indicated (**D**) FRET^+^ and (**E**) FRET^neg^ B cell compartments (black lines) to LPS, double-stranded DNA (dsDNA), single-stranded DNA (ssDNA), keyhole limpet hemocyanin (KLH) and insulin. mGO53 was included as non-reactive control antibody (green lines), ED38 was included as highly polyreactive control antibody (red lines). **(F, G)** Autoreactivity determined by the FLOWMIST assay. (**F**) Summary of antibody screening for autoreactivity by flow cytometry. Fold MFI relative to the negative control mGO53 is plotted. Dotted line represents the cutoff for positive reactivity. Control antibodies were included for comparison (mGO53: non-reactive, ED38: polyreactive, 673/719 and its germline revertant gl1/5: anti-nuclear). Reactive antibodies were confirmed by at least two independent experiments. (**G**) All autoreactive monoclonal antibodies identified in (F) are shown. Flow cytometry histograms for each antibody (red) are overlaid with the non-reactive mGO53 control antibody (grey). Confocal microscopy and ImageStream analysis depict the autoreactive fluorescence pattern (green) and DAPI (red). 5µm scale bars are shown on the bottom. **(H)** Monoclonal antibody screening for autoreactivity against C57Bl/6 spleen, bone marrow, kidney and thyroid lysates by dot blot. The relative grey value for the dot area of each antibody is shown normalized to non-reactive mGO53 antibody (grey). Highly polyreactive ED38 (blue) is included as positive control. Horizontal lines indicate cutoff for positive reactivity. (**A-H**) Monoclonal antibodies are derived from two to three independent single-cell sorts for each compartment.

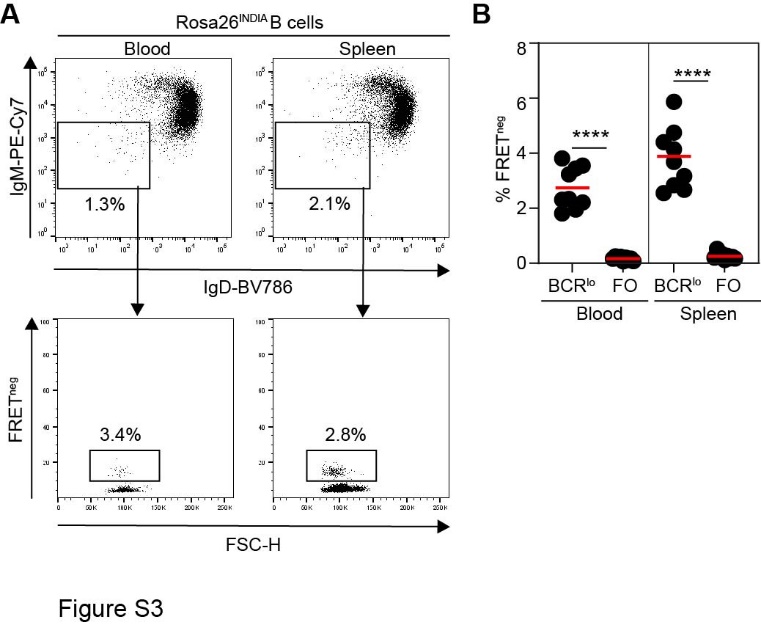

**Fig. S3. Death by neglect of T1 B cells.**

**(A, B)** Rosa26^INDIA^ mice were analyzed by flow cytometry. Blood and spleen B cells were gated CD19^+^mRuby2^+^DAPI^neg^Lineage(CD4, CD8α, F4/80, NK1.1, CD95)^neg^ and IgD^lo^IgM^lo^Igλ^neg^ (BCR^lo^) or IgD^hi^IgM^+^ (follicular, FO). **(A)** Representative dot plots (top) show gating of IgD^lo^IgM^lo^ B cells. Dot plots below show FSC-H and FRET loss of BCR^lo^ B cells. **(B)** Quantitation of FRET loss in BCR^lo^ and mature follicular (FO) B cells. Data are combined from three independent experiments each involving three animals (**** p<0.0001; unpaired student’s t-test).

**Auxiliary supplementary table legends**

**Table S1.** Summary of cloned BCR and antibody properties

**Table S2.** List of primers used for BCR cloning
